# Criticality and partial synchronization analysis in Wilson-Cowan and Jansen-Rit neural mass models

**DOI:** 10.1101/2023.10.03.560644

**Authors:** Sheida Kazemi, AmirAli Farokhniaee, Yousef Jamali

## Abstract

Synchronization is a phenomenon observed in neuronal networks involved in diverse brain activities. Neural mass models such as Wilson-Cowan (WC) and Jansen-Rit (JR) manifest synchronized states. Although they have been studied for decades, their ability to demonstrate second-order phase transition (SOPT) and criticality has not received enough attention, which serves as candidates for the development of healthy brain networks. In this study, two networks of coupled WC and JR nodes with small-world topologies were constructed and Kuramoto order parameter (KOP) was used to quantify the amount of synchronization. In addition, we investigated the presence of SOPT using the synchronization coefficient of variation. Both networks reached high synchrony by changing the coupling weight between their nodes. Moreover, they exhibited abrupt changes in the synchronization at certain values of the control parameter not necessarily related to a phase transition. While SOPT was observed only in JR model, neither WC nor JR model showed power-law behavior. Our study further investigated the global synchronization phenomenon that is known to exist in pathological brain states, such as seizure. JR model showed global synchronization, while WC model seemed to be more suitable in producing partially synchronized patterns.

**Corresponding author summary:** Yousef Jamali received his M.Sc. degree in physics from Sharif University, Tehran, Iran, in 2004, and his Ph.D. degree in physics from Sharif University, Tehran, Iran, in 2009. After finishing his Ph.D. thesis, in 2009, he got two years postdoctoral research associate position at the University of California at Berkeley, Department of Bioengineering. He is currently an Associate Professor in the Applied Mathematics Department (Biomathematics division), at Tarbiat Modares University, Tehran, Iran. His current research interests include the interdisciplinary area of the complex system especially on the modeling of brain. In fact, his interests are how the brain works at the critical state and how this complex system synchronized under different conditions.

## 1 Introduction

The dynamics of macroscopic and mesoscopic brain activity is a challenging topic. The large-scale model activities can be complex and include a range of behaviors such as spiking and oscillatory, resting-state, chaotic, and periodic or nonperiodic activities [1, 2]. The neural mass models which are based on the mesoscopic scale, describe the action of neural populations rather than the behavior of spiking neurons and have been used broadly in modeling some brain activities such as epilepsy [3], sleep [4], and human alpha rhythm (*∼* 8-*∼*12 Hz) [5]. These models depend on their output (voltage or rate) and are classified into two voltage- and activity-based models [6]. The first voltage-based model in respect of one excitatory and one inhibitory ensemble was built by Lopes da Silva [7]. As an extension of Lopes da Silva’s model, Zetterberg and colleagues included two excitatory and one inhibitory ensembles in their description of the cortex, an approach that Jansen and Rits adopted for the definition of their model [8, 9]. The activity-based models are also referred to as Wilson-Cowan (WC) equations [10], Amari equations [11], Cohen–Grossberg equations [12], and neural field equations [13]. These types of models exhibit a vast range of interesting and important dynamics such as Hopf bifurcation, which motivated researchers to establish their reduced version counterparts useful for dynamical systems studies [14, 15].

Neural activity synchronization is crucial to neural function and cognitive processes [16–19]. Synchronization of different single neuron classes to periodic external stimuli [20, 21] and coupled neurons [22–25] has been analyzed extensively, and can be measured both locally or over a global scale [26–29]. Synchronization, asynchronization, and partial synchronization are three cases of dynamical behavior in systems of coupled oscillators that manifest interesting and neurophysiologically relevant results. In previous studies of neural synchronization, most of the attention has been focused on global synchronization, which means how all nodes in a network behave in unison [30, 31]. However, the experimental results do not exhibit this property in a healthy brain. Some severe brain disorders, such as epilepsy, Parkinson’s, and essential tremor, may result from pathological synchronization [32–34]. However, partial synchronization within brain regions has been manifested for example in the macaque monkey’s visual system experimentally [35].

The conceptual appeal of the critical hypothesis is poising in a boundary between different types of dynamics that networks show the optimal performance of a cortical function [36–39]. Power-law behavior in various variables appears in critical dynamics. For example, the size and duration distributions of neuronal avalanches [36, 40] and EEG cascade [41, 42] obey a power-law. Moreover, long-range temporal correlations have been revealed in the amplitude envelopes of neural oscillations [43, 44].

The following study compares the dynamics of WC firing rate model and the voltage-based Jansen-Rit (JR) model. Historically, WC equations are regarded as a fundamental and classic model in computational neuroscience. According to predictions, the WC equations will still have great utility for several decades to come [45].JR is a minimal computational model of a small region of the cortex that belongs to a large group of models that are mathematically manifested by second-order linear differential equations. We simulated these two different types of neural mass models and investigated phase transition by taking into account the amount of synchronization as an order parameter. Then, their partial synchronization is discussed in the following sections. Our results showed that the coexistence between synchronization and asynchronization occurred simultaneously only in the WC model. Moreover, although we also observed continuous phase transition in JR model, neither of these models showed power-law behavior.

## 2 Methods

### 2.1 Wilson-Cowan model

The classical WC model describes the dynamics of firing rates among neural populations in the brain [10, 46]. The activity of each neural population is computed as the mean firing rate of its excitatory (*E*) and inhibitory (*I*) subpopulations by using mean-field approximation. The temporal evolution of the excitatory and inhibitory firing rates, *E*(*t*) and *I*(*t*), respectively, is governed by the following differential equations [10]:

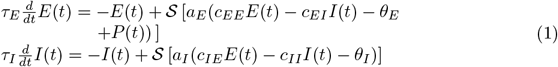

where *τ*_*E*_ (*τ*_*I*_) shows the time constant for excitatory (inhibitory) populations. The sigmoid function 𝒮 introduces the thresholds *θ*_*E*_ and *θ*_*I*_ corresponding to the maximum slope values and can be different for excitatory and inhibitory subpopulations. Moreover, the slopes of the sigmoids are given by *a*_*E*_ and *a*_*I*_ . 𝒮 is given as follows:

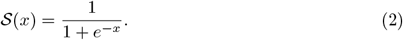

The synaptic weights are determined by the connectivity coefficients *c*_*EE*_, *c*_*EI*_, *c*_*IE*_, and *c*_*II*_ . *P* (*t*) represents the external stimuli acting on excitatory subpopulation in time t.

Depending on parameter settings, specifically on the selection of the external input P(t), the dynamics of this dynamical system ranges from fixed-point relaxations to limit cycle oscillations. Table 1 shows the values of parameters with their interpretations, which can produce oscillatory behavior in the WC model.

**Table 1.** Parameters in the WC model that shows the oscillatory behavior used in this study.

| Parameters | Interpretation | Value |
| --- | --- | --- |
| $\tau_E$ | Time constant for excitatory populations | 0.125 |
| $\tau_I$ | Time constant for inhibitory populations | 0.25 |
| $\theta_E$ | Maximum slope of sigmoid function for excitatory subpopulations | 2 |
| $\theta_I$ | Maximum slope of sigmoid function for inhibitory subpopulations | 8 |
| $a_E$ | The proportion of excitatory cells firing | 0.8 |
| $a_I$ | The proportion of inhibitory cells firing | 0.8 |
| $c_{EE}$ | Excitatory self-feedback synaptic weight | 8 |
| $c_{EI}$ | Synaptic weight from excitatory to inhibitory ensemble | 16 |
| $c_{IE}$ | Synaptic weight from inhibitory to excitatory ensemble | 8 |
| $c_{II}$ | Inhibitory self-feedback synaptic weight | 0.4 |

In a network with N nodes, i = 1,…, N, the WC equations are as follows:

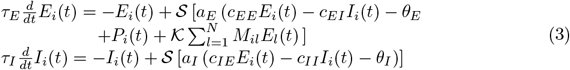

where *M*_*il*_ shows the network adjacency matrix and *𝒦* represents coupling coefficient between nodes.

An excitatory population interacts with an inhibitory population in each node (Fig. 1**(A)**). In a network, these nodes were linked through their connections with excitatory populations.

**Fig 1.**
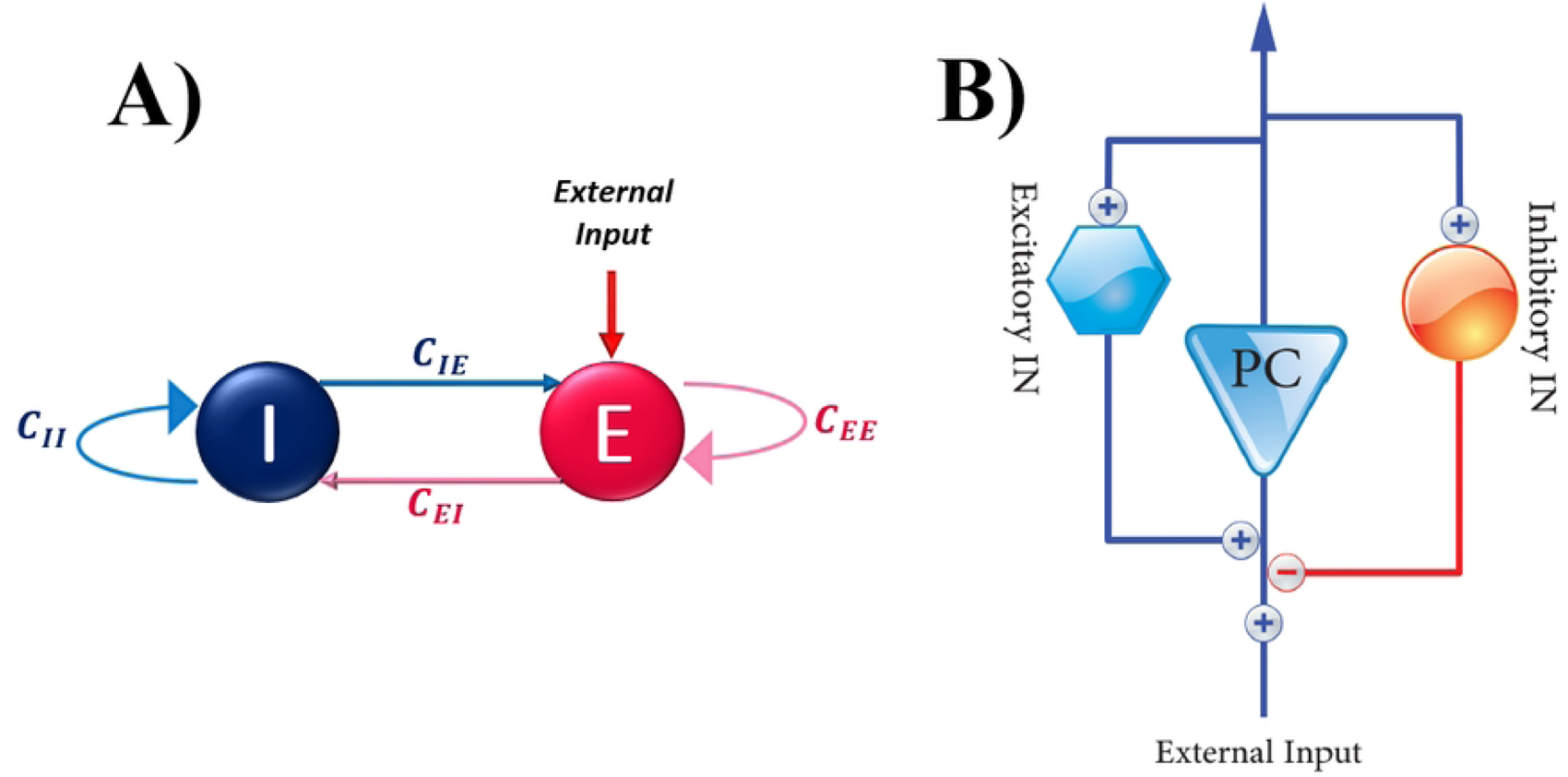
Schematic representation of WC (A) and JR (B). (A) Two ensembles of excitatory and inhibitory neurons. (B) A pyramidal cell (PC) with two excitatory and inhibitory interneurons in (**B**) interact with each other.

The appropriate choice of time constants *τ*_*E*_, *τ*_*I*_, and external inputs P(t) provided self-sustaining oscillations in the delta band frequency to match with the activity observed in [47].

### 2.2 Jansen-Rit model

JR model is based on the work of Lopes Da Silva [7]. A mathematical framework inspired by biology was developed to simulate spontaneous electrical activities of cortical columns, with an emphasis on alpha activity [8, 9].

In this model, pyramidal neurons receive input from excitatory and inhibitory interneurons in the same column as well as external input from other columns. This model can be written with six first-order differential equations as follows [8]:

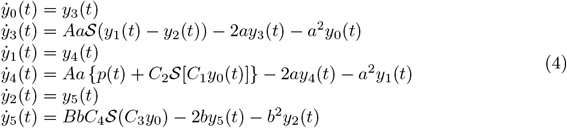

where *y*_*i*_ for *i* ∈ {0, 1, 2} represents the mean postsynaptic potentials of three neuronal populations (*y*_0_ for pyramidal, *y*_1_ for excitatory and *y*_2_ for inhibitory neurons). Their deviations are denoted by *y*_3_, *y*_4_, and *y*_5_, respectively. An external input is defined by the function *p*(*t*), which may be generated from external sources or by neighbouring neural populations.*S* is a sigmoidal function that transforms the average membrane potential of neurons into the mean firing rate of action potentials and is given by:

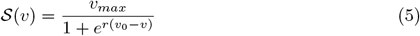

where *v*_*max*_ is the maximum firing rate of the neurons ensemble, *v*_0_ is potential at half of the maximum firing rate and *r* is the slope of the sigmoid at *v*_0_. In Table 2, all of the parameters are quantified. The output of this model is *y*_1_*−y*_2_, which represents the postsynaptic membrane potential of pyramidal neurons [48].

**Table 2.** Parameters in the JR model used in this study that are obtained experimentally.

| Parameters | Interpretation | Value |
| --- | --- | --- |
| $A$ | Excitatory PSP amplitude | $3.25mV$ |
| $B$ | Inhibitory PSP amplitude | $22mV$ |
| $1/a$ | Time constant of excitatory PSP | $a = 100s^{-1}$ |
| $1/b$ | Time constant of inhibitory PSP | $b = 50s^{-1}$ |
| $C1, C2$ | Average numbers of synapses between excitatory populations | $1 * C, 0.8 * C$ |
| $C3, C4$ | Average numbers of synapses between inhibitory populations | $0.25 * C$ |
| $C$ | Average numbers of synapses between neural populations | 135 |
| $v_{max}$ | Maximum firing rate | $5Hz$ |
| $v_0$ | Potential at half of maximum firing rate | $6mV$ |
| $r$ | Slope of sigmoid function at $v_0$ | $0.56mV^{-1}$ |

In a network with N nodes, i = 1,…, N, the JR equations are described by the following set of ordinary differential equations [49]:

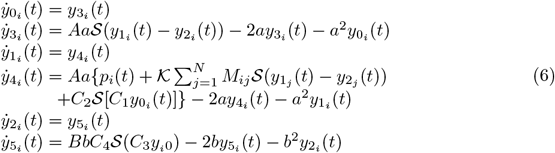

where *M*_*ij*_ is the adjacency matrix of the network and *𝒦* represents the coupling coefficient between nodes. A population of pyramidal neurons interacts with one excitatory and one inhibitory population of neurons, as schematically shown in Fig 1**(B)**. A network of nodes was formed by the connections between excitatory populations.

### 2.3 Network structure

Modeling biological, social, and many other real-world structures can be performed by networks of coupled dynamical systems, which include nodes (neurons or brain regions) and their physical connections. Although a regular or random connection topology is usually assumed in most cases, many real systems follow a different topology. Indeed, real networks are characterized by high clustering coefficients and small mean-shortest path lengths, which are observed in regular and random structures, respectively [50]. These networks with these two properties are known as small-world networks. The Watts-Strogatz topology is capable of generating graphs with small-world properties. A Watts-Strogatz topology is characterized by *N* (number of nodes in a ring), *K* (number of nearest neighbors of a node, K/2 on either side), and *p* (the probability of rewiring assigned to each edge). The appropriate choice of these three parameters has a substantial role to build a Watts-Strogatz model. It is best to choose K such that the resulting network is neither sparse nor fully connected. Moreover, the following equation should hold: *N* ≫ *K* ≫ ln *N* ≫ 1. *p* is the most challenging parameter to be estimated. Small-world properties are found in networks formed by 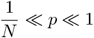.

### 2.4 Kuramoto order parameter

The Kuramoto model consists of interacting oscillators that each of them is considered to have its own intrinsic natural frequency *ω*, and each is coupled equally to all other oscillators. This model is most commonly governed by the following equations:

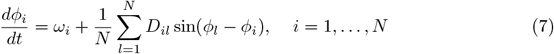

where *ϕ*_*i*_ is instantaneous phase of the *i*’s node, *D*_*il*_ is the phase coupling matrix, and *N* represents the number of nodes.

One way to recognize the degree of global synchronization in a network of coupled ensembles of identical oscillators is via the so-called Kuramoto order parameter (KOP). The value obtained with this criterion is equal to:

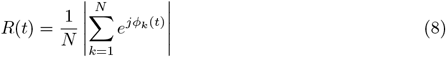

where *ϕ*_*k*_(*t*) followed the dynamics in equation (7) and 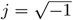. *R* gives a value range in [0, 1] with 0 representing no phase synchrony (asynchronous state) and *R* = 1 when full synchronization occurs. Notably, the phase calculated in the KOP can be the result of the Hilbert transform of the oscillating signals [51, 52].

### 2.5 Measure of criticality

The study of phase transitions has been investigated over many decades in a broad range of physical systems and it is one of the most active research areas in statistical mechanics [53]. A system with a second-order phase transition (SOPT) or continuous phase transition can sit at the transition point between two phases and separate an ordered and disordered one [54]. Therefore said to be on the edge of chaos. SOPT or criticality theory has sparked much interest over the years, as it has been demonstrated that critical behavior has many potential advantages such as maximal dynamical range, extensive information processing, and storage capabilities [55–57]. It is claimed that the healthy brain acts in a critical regime at a boundary between different types of dynamics [36–39]. It is a challenging question that how critical dynamics can be detected in neural models. There are some measures of criticality that can be assessed [58, 59]. In order for any marker of criticality to exist, first a critical point needs to be determined.

To investigate the SOPT, we compute the coefficient of variation (CV) of synchronization value against the control parameter (coupling coefficient). The peak in the CV during the continuous phase transition may be a marker of criticality [60].

It is documented that normal brain function is characterized by long-range temporal correlations (LRTCs) in the cortex [43, 44]. As well as this, the critical hypothesis is supported by the presence of LRTCs in neural oscillation amplitudes [61]. Veritably, long-range temporal correlations are crucial characteristic of criticality. Detrended Fluctuation Analysis (DFA) or mean auto-correlation methods are employed to investigate the temporal correlation structures of the signal. It is possible to detect LRTCs in a signal if its auto-correlation decays as a power-law with an exponent between −1 and 0 [62]. Generally, auto-correlation functions are very noisy in their tails, making exponent estimation difficult. In order to overcome these problems, DFA is an appropriate technique [44]. DFA determines the long-range temporal correlations in neuronal oscillation amplitude envelopes.

The DFA method can be explained as follows: first, a time series *X*(*t*) with a length of *N* and *t∈* 1, …, *N*, is divided into 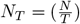 non-overlapping segments *X*_*i*_(*t*) with the same size *T* . A signal segment will be missed if (*N/T*) is not an integer. We divide signals from their end to overcome this problem. So, we have 2*N*_*T*_ segments. Then, the linear trend in segment *i* is taken out, and the fluctuations *F*_*i*_(2*N*_*T*_) corresponding to window length 2*N*_*T*_ are given as follows [63]:

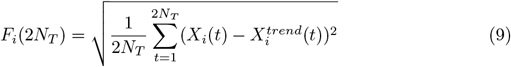

where 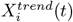 shows the linear trend in segment *i. F*_*i*_(2*N*_*T*_) is plotted against (2*N*_*T*_) in a logarithmic scale. In power-law relation, *F*_*i*_(2*N*_*T*_) *∼* (2*N*_*T*_)^*α*^ which *α* is determined as the slope of a straight line fit, and 0.5<*α*<1 represents long-range correlation.

### 2.6 Simulation

Several studies have used diffusion tensor imaging (DTI) data between region of interests (ROIs) of an atlas, which provided cortical and subcortical ROIs [28, 64, 65]. Following them, we construct a network of 80 identical coupled neural phase oscillators as nodes to better adapt to a real network. The connections between units follow the small-world topology (Watts-Strogatz network with a rewiring probability of 0.2). Each node is connected to 20 neighbors, ten on each side. Then, WC and JR dynamics were applied to each node individually. Different initial conditions for both models are adjusted to show oscillatory activity in each independent run. The length of this simulation is 200 s, and the time step is set to 10^*−*4^ s. For every coupling strength, each simulation is repeated twenty times. In this study, the network is analyzed by changing the global coupling strength *𝒦*.

## 3 Results

First, we consider the output series of each node and calculate the phase synchronization by KOP measure between them at each trial. Fig. 2 show the mean synchrony value against the coupling coefficient in JR (**A**) and WC (**B**) models. Vertical blue bars represent dispersions for twenty runs (standard deviations). According to the results in Fig. 2, depending on the coupling coefficient, different synchronization values arise from collective behaviors. For instance, complete synchronization on a network that can result in an oscillation state in which all oscillators behave the same way. Both models exhibited a transition from an unsynchronized to a highly synchronized state. This kind of behavior was detected in ensembles of coupled oscillators [47, 66–68]. A similar occurrence of transition from unsynchronized to full phase synchronized state is observed in [59].

**Fig 2.**
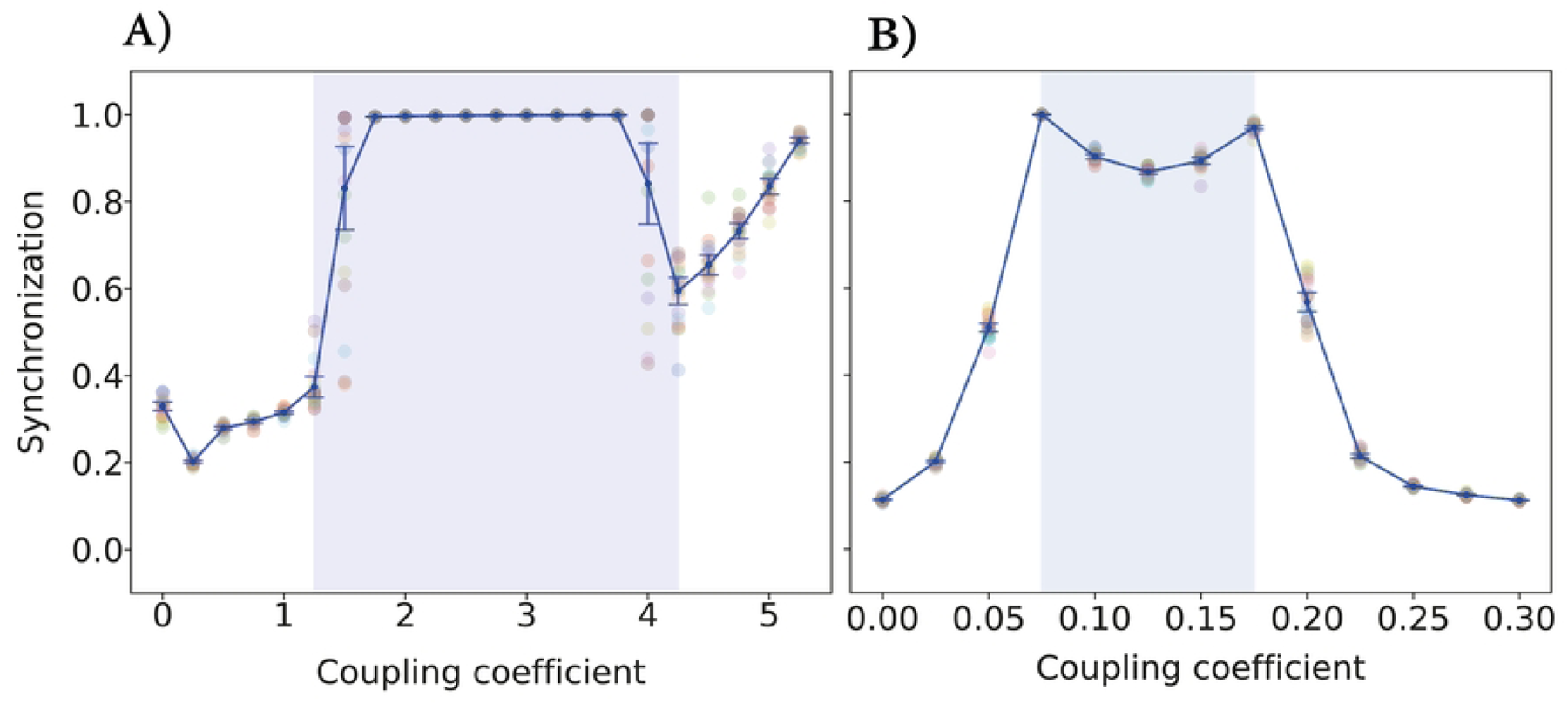
The KOP value according to the coupling coefficient in the JR (A) and WC (B) model. (A) At *𝒦* = 1.25 and 4.25, the dispersion of the synchronization value is high, and the area between them is related to the high synchrony regime. (B) At *𝒦* = 0.075 and 0.175, the blue highlighted regime shows an unexpected (decreasing in the synchrony value) pattern. The simulation runs twenty times on each coupling coefficient for these two models.

As shown in Fig. 2(**A**) increasing coupling up to 1.25 causes more synchronization. At a coupling coefficient equal to 1.25, the network’s synchronization has increased sharply, which can confirm a disorder in network function [69–71]. Our results indicated that in the [1.25, 4.25] regime of coupling strength, the synchronization values are high. We call this area a high synchrony regime and the coupling coefficient equal to 1.25 (4.25) is the starting (ending) point of high synchrony behavior. The coupling coefficient of 1.5 and 4 produces a large variation in the mean of the KOP, accompanied by a large variance, and can serve as the candidate for a phase transition point.

In Fig. 2(**B**), between *𝒦* = 0.075 to 0.175, the decrease in the synchrony value is seen. Indeed, Fig. 2(**B**) shows a local minimum in the KOP in which the increasing trend of the network by increasing coupling strength, the synchronization value decreases. It appears that this type of behavior also occurs in a network of Kuramoto model [72].

Now, to investigate the partial synchronization, we plot the colored output matrix of JR and WC network activity in Fig. 3 and 4, respectively. In a range of small coupling coefficients, the system activities are randomly distributed, related to incoherent states. In Fig. 3 the network shows the complete synchronization for 2 *< 𝒦 <* 4 which corresponding to coherent states. The nervous system in this condition is not in a healthy state and may lead to some brain disorder such as epileptic seizure activity [69, 73, 74]. Indeed, the system activity are coherent in space and time and shows the global synchronization.

**Fig 3.**
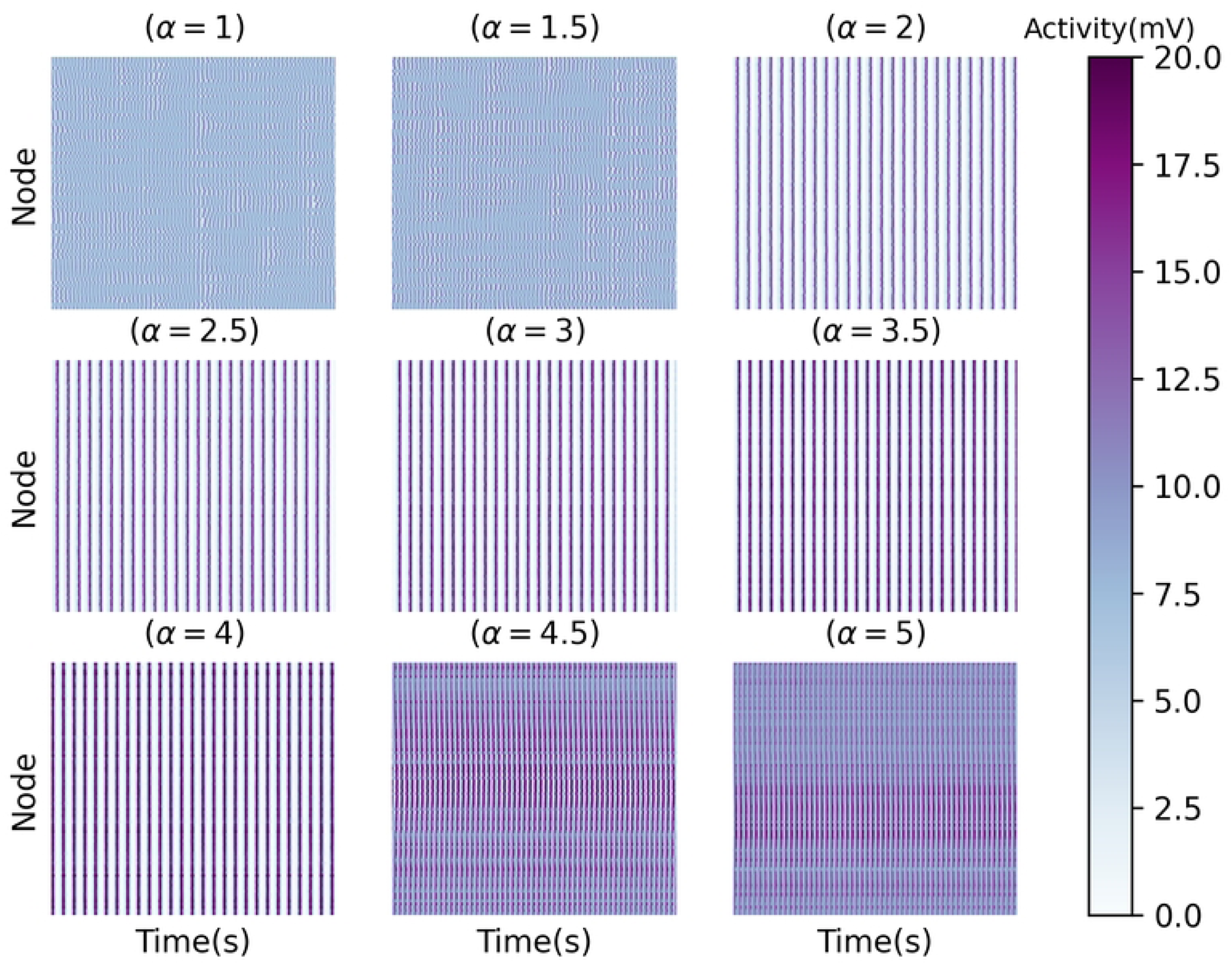
The colored output matrices during the last two seconds for different values of coupling coefficient in the JR model. In *𝒦* = 1.25 to 4.25, the output signal shows an ordered behavior.

**Fig 4.**
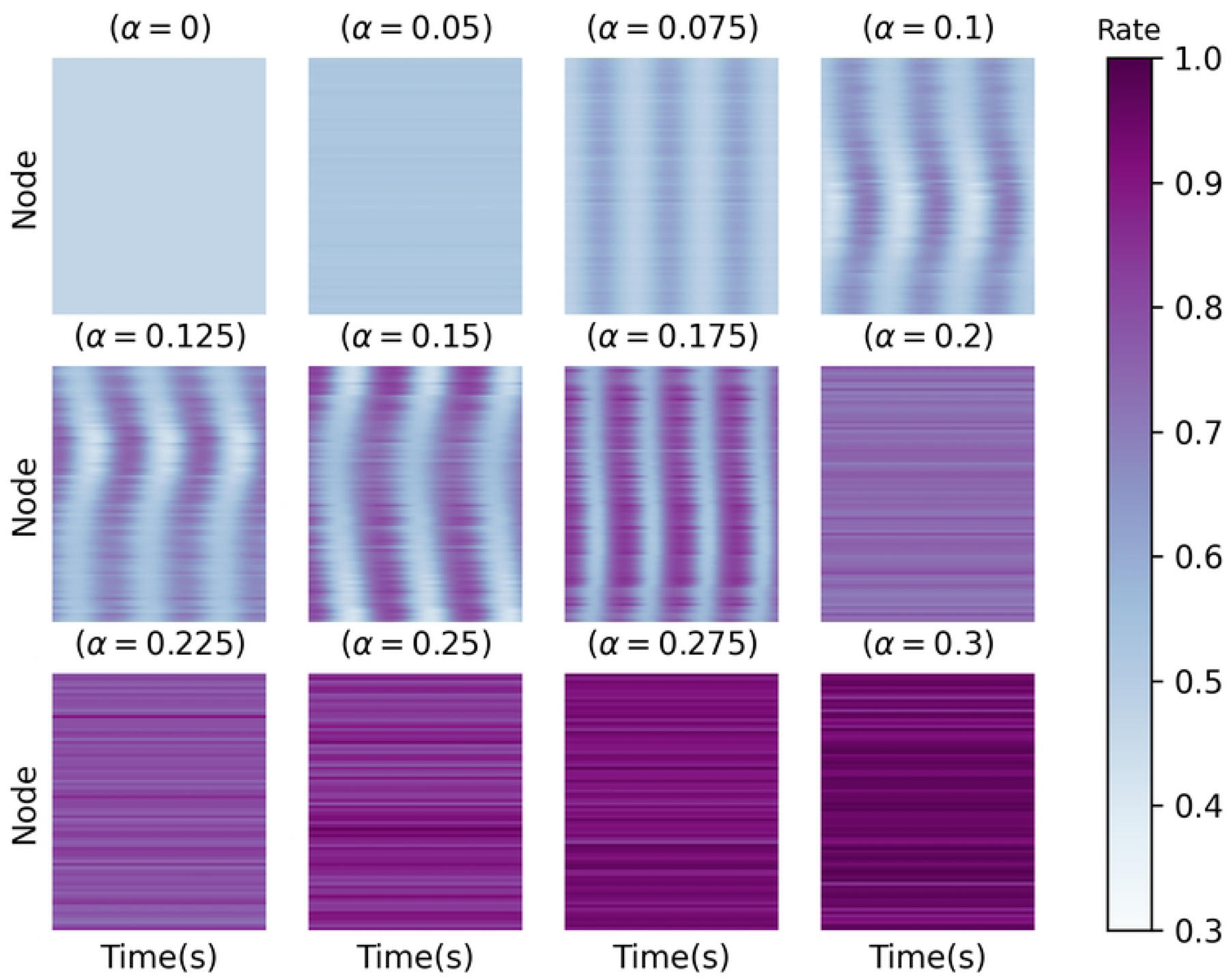
The colored output matrices during the last two seconds for different values of coupling coefficient in the WC model. In *𝒦* = 0.075 to 0.175, the output signal shows a slight time delay synchronization and the nodes are synchronized with a slight delay.

In Fig. 4, the colored activity matrix of the output signal shows a slight time delay synchronization corresponding to the blue region in Fig. 2(**B**) (0.075 *< 𝒦 <* 0.175). It means that the rate of the excitatory ensemble of neurons in WC model is coherent in time and incoherent in space. Indeed, the nodes are synchronized with delay to generate some patterns of partial synchronization that are evident only in this model and not in JR one.

A large number of neurologic disorders, including schizophrenia, Alzheimer’s disease, and Parkinson’s, may be associated with the coexistence of synchronization and asynchronization [16, 75, 76].

In order to analyze the SOPT, we compute the CV of Fig. 2 against the control parameter (coupling coefficient) in Fig. 5. Two peaks in the CV during the continuous phase transition (at *𝒦* =1.25, 4.25) are markers of criticality (Fig. 5(**A**)) in JR model. These points of maxima of this curve correspond to the value of the transition points in Fig. 2(**A**). Fig. 5(**B**) does not show any peak, which means WC model do not exhibit criticality.

**Fig 5.**
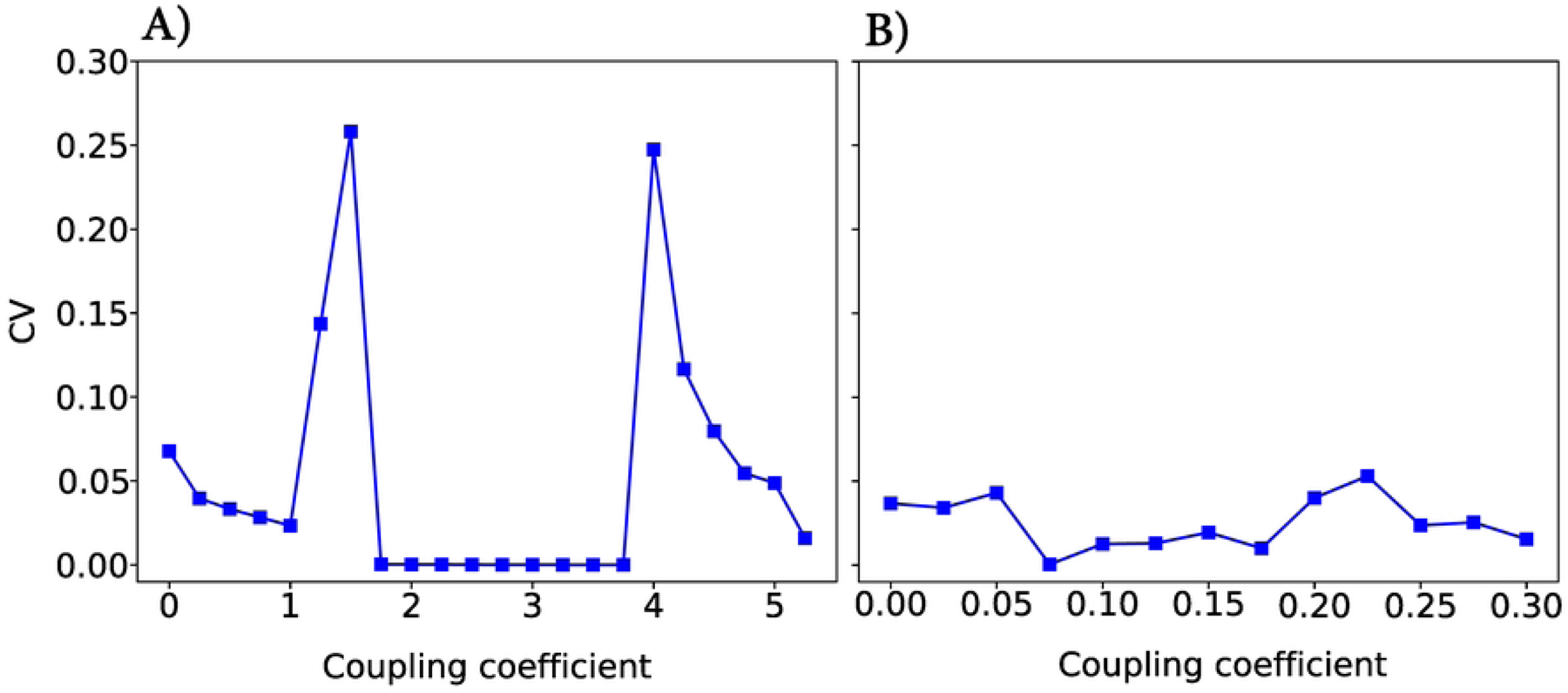
CV vs. coupling coefficient in (A) JR and (B) WC. The maxima in (A) correspond to transition points observed in Fig. 2.

Now, we examine LRTC in these two models. LRTCs in the amplitude of neural oscillations supports the critical dynamics of biological neuronal networks. Fig. 6 shows an arbitrary realization of the applied DFA technique in an absolute signal Hilbert transformation with no overlapping. Based on these results, it appears that the linear fit does not fit the data in either case. Hence, LRTCs are not present at these points. A piece-wise linear function (purple line) is suitable for fitting data which confirms the deviation from linearity. Similarly, the presence of LRTCs has not been observed in other coupling coefficients, so this feature is not apparent in either of these models with defined parameters. These results confirm previous findings [28, 59].

**Fig 6.**
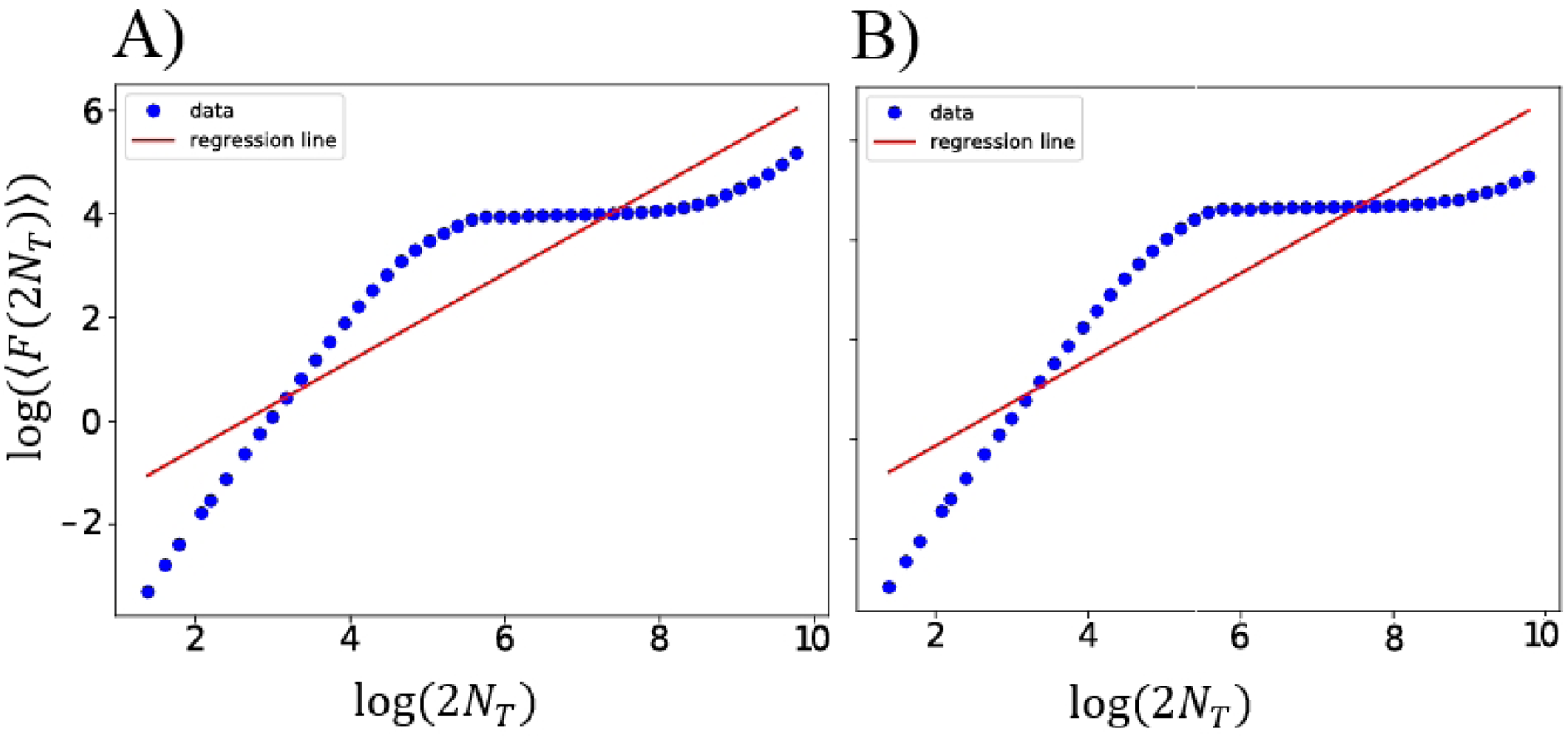
The fluctuation plot for an arbitrary value in JR (A) and WC (B) models in the log–log plot. The values on the x-axis are in seconds on logarithmic scales based on segment sizes, and the y-axis shows the standard deviation mean of all sized segments. This trend of data is piecewise linear, and the linear fit is not suitable for them. LRTCs do not exist at these points.

## 4 Discussion and Conclusion

In this study, we compared the dynamics of two seminal neural mass models, representing the integration of many neurons within a population. In the past few decades, neural mass models have contributed significantly to our understanding of meso- and macroscale dynamics of neuronal populations [47, 49, 66]. A wide range of neural phenomena has been modeled using them, such as alpha band oscillations [77], sleep rhythms [4], evoked potentials [8], and mechanisms of brain rhythms in disease [5, 78].

We investigated phase synchronization and criticality in two voltage-based (JR) and rate-based (WC) neural mass models. The two models were utilized in the same structural network representing the same topology. We observed two profound phenomena in brain activity. First, the global synchronicity of these networks was calculated by considering the KOP. The phase synchronization analysis of WC and JR neural mass models revealed that they shared a common feature. There was a transition from low to full synchronization in both models, which is not indicative of a healthy system, however, JR was the only model that showed a phase transition. This is likely because, in the WC model, the dynamics of a mass is expressed by a two-dimensional system. Indeed, WC model based on the equilibrium of both excitatory and inhibitory inputs shows only the fixed-points and oscillatory activities and may not describe the biophysical details of brain activity [10]. This model has revealed the presence of Hopf bifurcations [79]. But JR dynamics is described by a set of six ordinary differential equations and can differentiate between different brain waves. A single JR mass can produce different types of activities including epileptic spike-like activity and alpha-like ones that do not appear in WC model. Moreover, the detailed bifurcation analysis of JR has shown an impressively rich bifurcation diagram [78].

Another measure that we studied in these two networks is partial synchronization. Partial synchronization in neural models have attracted the attention of researchers. The emergence of different patterns in partial synchronization supports both healthy and disordered states in brain functions. In this work, we observed that WC model could not show the second-phase transition, however, does not exclude complex dynamics. This model generally synchronizes its nodes with a small time delay. Indeed, two synchronous and asynchronous behaviors was appeared simultaneously in this network, which was not seen in JR model.

Moreover, we investigated power-law behavior. Our results showed that even though JR model demonstrated a phase transition under the defined conditions, none of these models were capable of exhibiting power-law behavior and consequently criticality. Indeed, the occurrence of a continuous phase transition at a critical point is a necessary but not sufficient condition. Critical points exhibit clear characteristics and invariance of scale behavior. These phase transition points do not demonstrate scaling invariance, therefore they are not critical.

We are left with the question of whether any of these models can be modified so that they exhibit power-law and partial synchronization. Additionally, discovering the optimal neural mass model for investigating brain dynamics can be challenging. This question arises from the nature of the specific features selected to observe these findings. We also performed power spectrum analysis, bifurcation analysis, and entropy analysis of both networks, which were not included in the results section because they did not affect our understandings from the phase transition analysis. Indeed, it is a key to search for features that show a higher level of power analysis. As a final note, it is unclear how a different type of partial synchronization, such as cluster synchrony appear in these types of networks, which can be another research subject for future studies.

## Acknowledgment

The authors would like to thank the anonymous reviewers for their valuable comments and suggestions.

## References

1. Byrne Á, O’Dea RD, Forrester M, Ross J, Coombes S. Next-generation neural mass and field modeling. J Neurophysiol. 2020 Feb 1;123(2):726–742. doi: 10.1152/jn.00406.2019. Epub 2019 Nov 27.

2. Bick C, Goodfellow M, Laing CR, Martens EA. Understanding the dynamics of biological and neural oscillator networks through exact mean-field reductions: a review. J Math Neurosci. 2020 May 27;10(1):9. doi: 10.1186/s13408-020-00086-9.

3. Wendling F, Bartolomei F, Bellanger JJ, Chauvel P. Epileptic fast activity can be explained by a model of impaired GABAergic dendritic inhibition. Eur J Neurosci. 2002 May;15(9):1499–508. doi: 10.1046/j.1460-9568.2002.01985.x.

4. Phillips AJ, Robinson PA. A quantitative model of sleep-wake dynamics based on the physiology of the brainstem ascending arousal system. J Biol Rhythms. 2007 Apr;22(2):167–79. doi: 10.1177/0748730406297512.

5. Freyer F, Roberts JA, Becker R, Robinson PA, Ritter P, Breakspear M. Biophysical mechanisms of multistability in resting-state cortical rhythms. J Neurosci. 2011 Apr 27;31(17):6353–61. doi: 10.1523/JNEUROSCI.6693-10.2011.

6. Ermentrout, B. Neural networks as spatio-temporal pattern-forming systems. Rep. Prog. Phys. 1998 Feb 2;4(61):353–430. doi: 10.1088/0034-4885/61/4/002.

7. Lopes da Silva FH, Hoeks A, Smits H, Zetterberg LH. Model of brain rhythmic activity: the alpha-rhythm of the thalamus. Kybernetik. 1974 May 31;15(1):27–37. doi: 10.1007/BF00270757.

8. Jansen BH, Zouridakis G, Brandt ME. A neurophysiologically-based mathematical model of flash visual evoked potentials. Biol Cybern. 1993;68(3):275–83. doi: 10.1007/BF00224863.

9. Jansen BH, Rit VG. Electroencephalogram and visual evoked potential generation in a mathematical model of coupled cortical columns. Biol Cybern. 1995 Sep;73(4):357–66. doi: 10.1007/BF00199471.

10. Wilson HR, Cowan JD. Excitatory and inhibitory interactions in localized populations of model neurons. Biophys J. 1972 Jan;12(1):1–24. doi: 10.1016/S0006-3495(72)86068-5.

11. Jaeger, D., Jung, R. (Eds.). Encyclopedia of computational neuroscience. New York, NY: Springer New York.; 2015.

12. Lu W, Chen T. New conditions on global stability of Cohen-Grossberg neural networks. Neural Comput. 2003 May;15(5):1173–89. doi: 10.1162/089976603765202703.

13. Coombes, S. Waves, bumps, and patterns in neural field theories. Biol Cybern. 2005 Aug;93(2):91–108. doi: 10.1007/s00422-005-0574-y.

14. Hoppensteadt, F.C. Weakly connected neural networks. Springer Science & Business Media.; 2005.

15. Farokhniaee, A., Almonte, F. V., Yelin, S., Large, E. W. Entrainment of weakly coupled canonical oscillators with applications in gradient frequency neural networks using approximating analytical methods. Mathematics. 2020 Aug;8(8):1312. 91-108. doi: 10.3390/math8081312.

16. Uhlhaas PJ, Singer W. Neural synchrony in brain disorders: relevance for cognitive dysfunctions and pathophysiology. Neuron. 2006 Oct 5;52(1):155–68. doi: 10.1016/j.neuron.2006.09.020.

17. Jutras MJ, Fries P, Buffalo EA. Gamma-band synchronization in the macaque hippocampus and memory formation. J Neurosci. 2009 Oct 7;29(40):12521–31. doi: 10.1523/JNEUROSCI.0640-09.2009.

18. Reinhart RMG. Synchronizing neural rhythms. Science. 2022 Aug 5;377(6606):588–589. doi: 10.1126/science.add4834.

19. Lowet E, De Weerd P, Roberts MJ, Hadjipapas A. Tuning Neural Synchronization: The Role of Variable Oscillation Frequencies in Neural Circuits. Front Syst Neurosci. 2022 Jul 8;16:908665. doi: 10.3389/fnsys.2022.908665.

20. Farokhniaee A, Large EW. Mode-locking behavior of Izhikevich neurons under periodic external forcing. Phys Rev E. 2017 Jun;95(6-1):062414. doi: 10.1103/PhysRevE.95.062414.

21. Farokhniaee A, Large EW. Mode-locking behavior of Izhikevich neurons under periodic external forcing. BMC Neuroscience. 2015 Jun;16(Supp 1):P140. doi: 10.1186/1471-2202-16-S1-P140.

22. Boccaletti, S., Kurths, J., Osipov, G., Valladares, D. L., Zhou, C. S. The synchronization of chaotic systems. Phys. Rep. 2002 Jun;366(1-2):1–101. doi: 10.1016/S0370-1573(02)00137-0.

23. Kumar R, Bilal S, Ramaswamy R. Synchronization properties of coupled chaotic neurons: The role of random shared input. Chaos. 2016 Jun;26(6):063118. doi: 10.1063/1.4954377.

24. Protachevicz PR, Hansen M, Iarosz KC, Caldas IL, Batista AM, Kurths J. Emergence of neuronal synchronisation in coupled areas. Front Comput Neurosci. 2021 Apr 22;15:663408. doi: 10.3389/fncom.2021.663408.

25. Cooray GK, Rosch RE, Friston KJ. Global dynamics of neural mass models. PLoS Comput Biol. 2023 Feb 10;19(2):e1010915. doi: 10.1371/journal.pcbi.1010915.

26. Jiruska P, de Curtis M, Jefferys JG, Schevon CA, Schiff SJ, Schindler K. Synchronization and desynchronization in epilepsy: controversies and hypotheses. J Physiol. 2013 Feb 15;591(4):787–97. doi: 10.1113/jphysiol.2012.239590.

27. Morales AJ, Vavilala V, Benito RM, Bar-Yam Y. Global patterns of synchronization in human communications. J R Soc Interface. 2017 Mar;14(128):20161048. doi: 10.1098/rsif.2016.1048.

28. Daffertshofer A, Ton R, Pietras B, Kringelbach ML, Deco G. Scale-freeness or partial synchronization in neural mass phase oscillator networks: Pick one of two?. Neuroimage. 2018 Oct 15;180(Pt B):428–441. doi: 10.1016/j.neuroimage.2018.03.070.

29. Kori H, Kiss IZ, Jain S, Hudson JL. Partial synchronization of relaxation oscillators with repulsive coupling in autocatalytic integrate-and-fire model and electrochemical experiments. Chaos. 2018 Apr;28(4):045111. doi: 10.1063/1.5022497.

30. Lee HY, Jung KI, Yoo WK, Ohn SH. Global synchronization index as an indicator for tracking cognitive function changes in a traumatic brain injury patient: a case report. Ann Rehabil Med. 2019 Feb;43(1):106–110. doi: 10.5535/arm.2019.43.1.106.

31. Ziaeemehr A, Zarei M, Sheshbolouki A. Emergence of global synchronization in directed excitatory networks of type I neurons. Sci Rep. 2020 Feb 24;10(1):3306. doi: 10.1038/s41598-020-60205-0.

32. Nini A, Feingold A, Slovin H, Bergman H. Neurons in the globus pallidus do not show correlated activity in the normal monkey, but phase-locked oscillations appear in the MPTP model of parkinsonism. J Neurophysiol. 1995 Oct;74(4):1800–5. doi: 10.1152/jn.1995.74.4.1800.

33. Bergman H, Deuschl G. Pathophysiology of Parkinson’s disease: from clinical neurology to basic neuroscience and back. Mov Disord. 2002;17 Suppl 3:S28–40. doi: 10.1002/mds.10140.

34. Tass PA, Silchenko AN, Hauptmann C, Barnikol UB, Speckmann EJ. Long-lasting desynchronization in rat hippocampal slice induced by coordinated reset stimulation. Phys Rev E Stat Nonlin Soft Matter Phys. 2009 Jul;80(1 Pt 1):011902. doi: 10.1103/PhysRevE.80.011902.

35. Drebitz E, Haag M, Grothe I, Mandon S, Kreiter AK. Attention configures synchronization within local neuronal networks for processing of the behaviorally relevant stimulus. Front Neural Circuits. 2018 Aug 29;12:71. doi: 10.3389/fncir.2018.00071.

36. Beggs JM, Plenz D. Neuronal avalanches in neocortical circuits. J Neurosci. 2003 Dec 3;23(35):11167–77. doi: 10.1523/JNEUROSCI.23-35-11167.2003.

37. Kitzbichler MG, Smith ML, Christensen SR, Bullmore E. Broadband criticality of human brain network synchronization. PLoS Comput Biol. 2009 Mar;5(3):e1000314. doi: 10.1371/journal.pcbi.1000314.

38. Tetzlaff C, Okujeni S, Egert U, Wörgötter F, Butz M. Self-organized criticality in developing neuronal networks. PLoS Comput Biol. 2010 Dec 2;6(12):e1001013. doi: 10.1371/journal.pcbi.1001013.

39. Meisel C, Storch A, Hallmeyer-Elgner S, Bullmore E, Gross T. Failure of adaptive self-organized criticality during epileptic seizure attacks. PLoS Comput Biol. 2012 Jan;8(1):e1002312. doi: 10.1371/journal.pcbi.1002312.

40. Beggs JM, Plenz D. Neuronal avalanches are diverse and precise activity patterns that are stable for many hours in cortical slice cultures. J Neurosci. 2004 Jun 2;24(22):5216–29. doi: 10.1523/JNEUROSCI.0540-04.2004.

41. Miskovic V, MacDonald KJ, Rhodes LJ, Cote KA. Changes in EEG multiscale entropy and power-law frequency scaling during the human sleep cycle. Hum Brain Mapp. 2019 Feb 1;40(2):538–551. doi: 10.1002/hbm.24393.

42. Jannesari M, Saeedi A, Zare M, Ortiz-Mantilla S, Plenz D, Benasich AA. Stability of neuronal avalanches and long-range temporal correlations during the first year of life in human infants. Brain Struct Funct. 2020 Apr;225(3):1169–1183. doi: 10.1007/s00429-019-02014-4.

43. Linkenkaer-Hansen K, Nikouline VV, Palva JM, Ilmoniemi RJ. Long-range temporal correlations and scaling behavior in human brain oscillations. J Neurosci. 2001 Feb 15;21(4):1370–7. doi: 10.1523/JNEUROSCI.21-04-01370.2001.

44. Hardstone R, Poil SS, Schiavone G, Jansen R, Nikulin VV, Mansvelder HD, Linkenkaer-Hansen K. Detrended fluctuation analysis: a scale-free view on neuronal oscillations. Front Physiol. 2012 Nov 30;3:450. doi: 10.3389/fphys.2012.00450.

45. Chow CC, Karimipanah Y. Before and beyond the Wilson–Cowan equations. J Neurophysiol. 2020 May 1;123(5):1645–1656. doi: 10.1152/jn.00404.2019.

46. Wilson HR, Cowan JD. A mathematical theory of the functional dynamics of cortical and thalamic nervous tissue. Kybernetik. 1973 Sep;13(2):55–80. doi: 10.1007/BF00288786.

47. Nazemi PS, Jamali Y. On the influence of structural connectivity on the correlation patterns and network synchronization. Front Comput Neurosci. 2019 Jan 8;12:105. doi: 10.3389/fncom.2018.00105.

48. Kandel, E. R., Schwartz, J. H., Jessell, T. M., Siegelbaum, S., Hudspeth, A. J., Mack, S. (Eds.). Principles of neural science. New York: McGraw-hill. ; 2000.

49. Forrester M, Crofts JJ, Sotiropoulos SN, Coombes S, O’Dea RD. The role of node dynamics in shaping emergent functional connectivity patterns in the brain. Netw Neurosci. 2020 May 1;4(2):467–483. doi: 10.1162/netna00130.

50. Watts DJ, Strogatz SH. Collective dynamics of ‘small-world’ networks. Nature. 1998 Jun 4;393(6684):440–2. doi: 10.1038/30918.

51. Kuramoto, Y. International symposium on mathematical problems in theoretical physics. Lect. Notes Phys. 1975 Jan 30;420. doi: 10.1007/BFb0013294.

52. Strogatz SH. From Kuramoto to Crawford: exploring the onset of synchronization in populations of coupled oscillators. Lect. Notes Phys. 2000 Sep 143(1-4):1–20. doi: 10.1016/S0167-2789(00)00094-4.

53. Papon, P., Leblond, J., Meijer, P. H. Physics of Phase Transitions. Berlin Heidelberg, Germany: Springer-Verlag.; 002.

54. Ojovan, M. I. Ordering and structural changes at the glass–liquid transition. J. Non-Cryst. 2013 Dec 382:79–86. doi: 10.1016/j.jnoncrysol.2013.10.016.

55. Kauffman SA, Johnsen S. Coevolution to the edge of chaos: Coupled fitness landscapes, poised states, and coevolutionary avalanches. J Theor Biol. 1991 Apr 21;149(4):467–505. doi: 10.1016/s0022-5193(05)80094-3.

56. O’Byrne J, Jerbi K. Are biological systems poised at criticality?. Trends Neurosci. 2022 Nov;45(11):820–837. doi: 10.1016/j.tins.2022.08.007.

57. Nykter M, Price ND, Larjo A, Aho T, Kauffman SA, Yli-Harja O, Shmulevich I. Critical networks exhibit maximal information diversity in structure-dynamics relationships. Phys Rev Lett. 2008 Feb 8;100(5):058702. doi: 10.1103/PhysRevLett.100.058702.

58. Hesse J, Gross T. Self-organized criticality as a fundamental property of neural systems. Front Syst Neurosci. 2014 Sep 23;8:166. doi: 10.3389/fnsys.2014.00166.

59. Kazemi S, Jamali Y. Phase synchronization and measure of criticality in a network of neural mass models. Sci Rep. 2022 Jan 25;12(1):1319. doi: 10.1038/s41598-022-05285-w.

60. di Santo S, Villegas P, Burioni R, Muñoz MA. Landau–Ginzburg theory of cortex dynamics: Scale-free avalanches emerge at the edge of synchronization. Proc Natl Acad Sci U S A. 2018 Feb 13;115(7):E1356–E1365. doi: 10.1073/pnas.1712989115.

61. Botcharova M, Farmer SF, Berthouze L. Markers of criticality in phase synchronization. Front Syst Neurosci. 2014 Sep 24;8:176. doi: 10.3389/fnsys.2014.00176.

62. Talkner P, Weber RO. Power spectrum and detrended fluctuation analysis: Application to daily temperatures. Phys Rev E Stat Phys Plasmas Fluids Relat Interdiscip Topics. 2000 Jul;62(1 Pt A):150–60. doi: 10.1103/physreve.62.150.

63. Peng CK, Havlin S, Stanley HE, Goldberger AL. Quantification of scaling exponents and crossover phenomena in nonstationary heartbeat time series. Chaos. 1995;5(1):82–7. doi: 10.1063/1.166141.

64. Van Essen DC, Smith SM, Barch DM, Behrens TE, Yacoub E, Ugurbil K; WU-Minn HCP Consortium. The WU-Minn human connectome project: an overview. Neuroimage. 2013 Oct 15;80:62–79. doi: 10.1016/j.neuroimage.2013.05.041.

65. Kashyap A, Keilholz S. Dynamic properties of simulated brain network models and empirical resting-state data. Netw Neurosci. 2019 Feb 1;3(2):405–426. doi: 10.1162/netna00070.

66. Daffertshofer A, van Wijk BC. On the influence of amplitude on the connectivity between phases. Front Neuroinform. 2011 Jul 15;5:6. doi: 10.3389/fninf.2011.00006.

67. Budzinski RC, Boaretto BRR, Prado TL, Viana RL, Lopes SR. Nonstationary transition to phase synchronization of neural networks induced by the coupling architecture. Chaos. 2019 Dec;29(12):123132. doi: 10.1063/1.5128495.

68. Budzinski, R. C., Boaretto, B. R. R., Prado, T. L., Lopes, S. R. Investigation of details in the transition to synchronization in complex networks by using recurrence analysis. Math. Comput. Appl. 2019 Apr;24(2):42. doi: 10.3390/mca24020042.

69. Sohanian Haghighi H, Markazi AHD. A new description of epileptic seizures based on dynamic analysis of a thalamocortical model. Sci Rep. 2017 Oct 19;7(1):13615. doi: 10.1038/s41598-017-13126-4.

70. Paula CAR, Reategui C, Costa BKS, da Fonseca CQ, da Silva L, Morya E, Brasil FL. High-frequency EEG variations in children with autism spectrum disorder during human faces visualization. Biomed Res Int. 2017;2017:3591914. doi: 10.1155/2017/3591914.

71. Igberaese, A. E., Tcheslavski, G. V. EEG power spectrum as a biomarker of autism: a pilot study. IJEH. 2018 May; 10(4):275–286. doi: 10.1504/IJEH.2018.101446.

72. Frolov N, Maksimenko V, Majhi S, Rakshit S, Ghosh D, Hramov A. Chimera-like behavior in a heterogeneous Kuramoto model: The interplay between attractive and repulsive coupling. Chaos. 2020 Aug;30(8):081102. doi: 10.1063/5.0019200.

73. Breakspear M, Roberts JA, Terry JR, Rodrigues S, Mahant N, Robinson PA. A unifying explanation of primary generalized seizures through nonlinear brain modeling and bifurcation analysis. Cereb Cortex. 2006 Sep;16(9):1296–313. doi: 10.1093/cercor/bhj072.

74. Maturana MI, Meisel C, Dell K, Karoly PJ, D’Souza W, Grayden DB, Burkitt AN, Jiruska P, Kudlacek J, Hlinka J, Cook MJ, Kuhlmann L, Freestone DR. Critical slowing down as a biomarker for seizure susceptibility. Nat Commun. 2020 May 1;11(1):2172. doi: 10.1038/s41467-020-15908-3.

75. Protachevicz PR, Borges FS, Lameu EL, Ji P, Iarosz KC, Kihara AH, Caldas IL, Szezech JD Jr, Baptista MS, Macau EEN, Antonopoulos CG, Batista AM, Kurths J. Bistable firing pattern in a neural network model. Front Comput Neurosci. 2019 Apr 5;13:19. doi: 10.3389/fncom.2019.00019.

76. Coninck, J. C., Ferrari, F. A., Reis, A. S., Iarosz, K. C., Caldas, I. L., Batista, A. M., Viana, R. L. Network properties of healthy and Alzheimer brains. Phys. A: Stat. Mech. 2020 June 547:124475. doi: 10.1016/j.physa.2020.124475.

77. Grimbert F, Faugeras O. Bifurcation analysis of Jansen’s neural mass model. Neural Comput. 2006 Dec;18(12):3052–68. doi: 10.1162/neco.2006.18.12.3052.

78. Touboul J, Wendling F, Chauvel P, Faugeras O. Neural mass activity, bifurcations, and epilepsy. Neural Comput. 2011 Dec;23(12):3232–86. doi: 10.1162/NECOa00206.

79. Li X, Li Z, Yang W, Wu Z, Wang J. Bidirectionally regulating gamma oscillations in wilson-cowan model by self-feedback loops: A computational study. Front Syst Neurosci. 2022 Feb 21;16:723237. doi: 10.3389/fnsys.2022.723237.

